# Accurate analysis of genuine CRISPR editing events with ampliCan

**DOI:** 10.1101/249474

**Authors:** Kornel Labun, Xiaoge Guo, Alejandro Chavez, George Church, James Gagnon, Eivind Valen

**Affiliations:** Department of Informatics/Computational Biology Unit; University of Bergen; Bergen, 5008; Norway; Wyss Institute for Biologically Inspired Engineering; Harvard University; Cambridge, MA 02115; USA; Department of Genetics; Harvard Medical School; Boston, Massachusetts, 02115; USA; Department of Pathology and Cell Biology; Columbia University; New York, NY 10032; USA; Department of Biology; University of Utah; Salt Lake City, UT 84112; USA; Sars International Centre for Marine Molecular Biology; University of Bergen; Bergen, 5008; Norway

**Author notes:** Correspondence: Eivind Valen.

## Abstract

We present ampliCan, an analysis tool for genome editing that unites highly precise quantification and visualization of genuine genome editing events. ampliCan features nuclease-optimized alignments, filtering of experimental artifacts, event-specific normalization, off-target read detection and quantifies insertions, deletions, HDR repair as well as targeted base editing. It is scalable to thousands of amplicon sequencing-based experiments from any genome editing experiment, including CRISPR. It enables automated integration of controls and accounts for biases at every step of the analysis. We benchmarked ampliCan on both real and simulated datasets against other leading tools, demonstrating that it outperformed all in the face of common confounding factors.

## Introduction

With the introduction of CRISPR (Jinek et al. 2012; Cong et al. 2013), researchers obtained an inexpensive and effective tool for targeted mutagenesis. Despite some limitations, CRISPR has been widely adopted in research settings and has made inroads into medical applications (Courtney et al. 2016). Successful genome editing relies on the ability to confidently identify induced mutations after repair through non-homologous end-joining (NHEJ) or homology directed repair (HDR). Insertions or deletions (indels) are often identified by sequencing the targeted loci and comparing the sequenced reads to a reference sequence. Deep sequencing has the advantage of both capturing the nature of the indel, readily identifying frameshift mutations or disrupted regulatory elements, and characterizing the heterogeneity of the introduced mutations in a population. This is of particular importance when the aim is allele-specific editing or the experiment can result in mosaicism.

The reliability of a sequencing-based approach is dependent on the processing and interpretation of the sequenced reads and is contingent on factors such as the inclusion of controls, the alignment algorithm and the filtering of experimental artifacts. To date, no tool considers and controls for the whole range of biases that can influence this interpretation and therefore, distorts the estimate of the mutation efficiency and leads to erroneous conclusions. Here we introduce a fully automated tool, ampliCan, designed to determine the true mutation frequencies of CRISPR experiments from high-throughput DNA amplicon sequencing. It scales to genome-wide experiments and can be used alone or integrated with the CHOPCHOP (Montague et al. 2014; Labun et al. 2016) guide RNA (gRNA) design tool.

## Results

### ampliCan accurately determines the true mutation efficiency

Estimation of the true mutation efficiency depends on multiple steps all subject to different biases (Lindsay et al. 2016). Following sequencing, reads have to be aligned to the correct reference, filtered for artifacts, and then the mutation efficiency has to be quantified and normalized (**Fig. 1A**). In most existing tools, many of the choices made during these steps are typically hidden from the user leading to potential misinterpretation of the data. These hidden steps can lead to widely different estimates of mutation efficiency (in up to 67% of all experiments) when run on data from real experiments (**Supplementary Note 3** and **Supplementary Fig. 3**). Furthermore, steps are frequently relegated to other tools that have not been optimized for CRISPR experiments. ampliCan instead implements a complete pipeline from alignment to interpretation and can therefore control for biases at every step.

**Fig 1.**
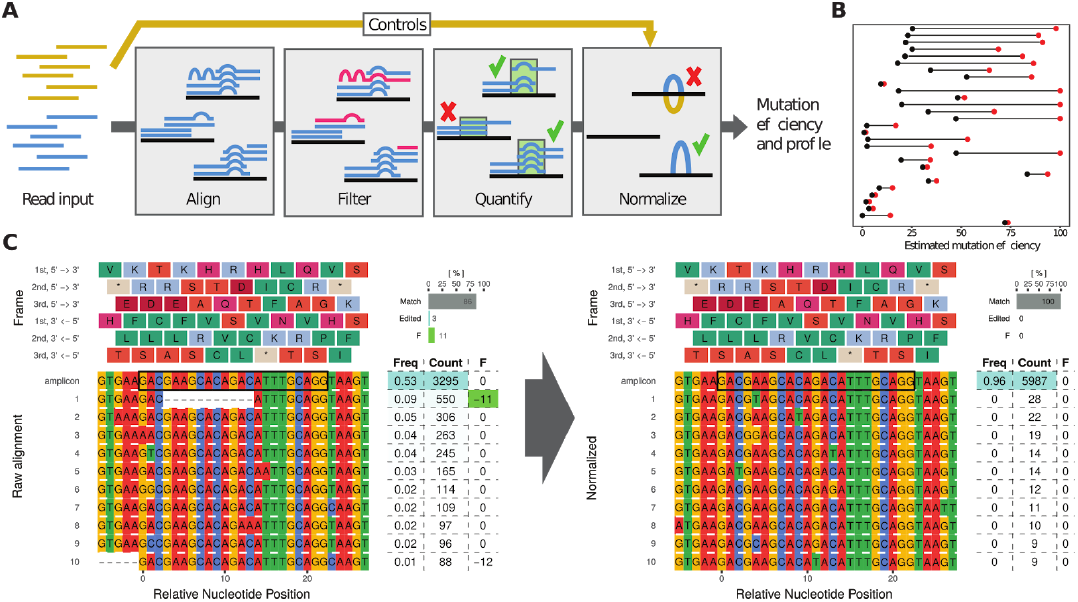
**A.** Estimation of mutation efficiency consists of multiple steps. At each of these steps biases can be introduced. Controls are processed identically to the main experiment and used for normalization. **B.** Overview of the change in estimated mutation efficiency on real CRISPR experiments when using controls that account for natural genetic variance in 29 experiments (mean change of 30%). Red dots show initial estimates based on unnormalized data, while black dots show the values after normalization. **C.** Alignment plot showing the top 10 most abundant reads in a real experiment. The table shows relative efficiency (Freq) of read, absolute number of reads (Count) and the summed size of the indel(s) (F), coloured green when inducing a frameshift. The bars (top right) shows the fraction of reads that contain no indels (Match), those having an indel without inducing frameshift (InDel) and frameshift inducing indels (F). The left panel shows the estimated mutation efficiency from raw reads, which is 14% (11% with frameshift, 3% without). The right panel shows the same genomic loci after normalization with controls resulting in an mutation efficiency of 0%. The deletion of 11bp in 9% of the reads could not be found in GRCz10.88 Ensembl Variation database and would in the absence of controls give the impression of a real editing event.

Despite being arguably the most important step in any experiment, the use of controls is frequently overlooked in CRISPR assays. Discrepancies between a reference genome and the genetic variation in an organism of interest often lead to false positives and the false impression that mutations have been introduced (Gagnon et al. 2014). While the use of controls is (in principle) possible with any tool, it commonly requires running the treated and control samples separately followed by a manual inspection and comparison of these. In ampliCan, controls are an integrated part of the pipeline and mutation frequencies are normalized and estimated automatically. ampliCan accomplishes this by normalizing at the event-level rather than the read-level. Any difference to the reference sequence (insertion, deletion or mismatch) that occurs in the controls above the level of noise is ignored when calculating mutation frequencies in the edited sample. This process is blind to the source of the event which may include genetic variance as well as experimental and sequencing artifacts. Because the normalization process does not remove any reads it also does not remove genuine editing events that may co-occur with a normalized event (see **Supplementary Note 1**). To assess the impact of controls we generated 112 CRISPR datasets and pooled them with data we previously generated in Gagnon et al. 2014 for a total of 263 experiments (Methods and **Supplementary Note 3**). These consisted of pools of CRISPR-injected zebrafish using wild type fish as control. This experimental setup presents a challenging task to pipelines since the genetic background may not be identical across all fish and the injected fish can be highly mosaic in their mutational outcomes. This benchmark revealed that accounting for the genetic background in the wild type fish reduced the estimated mutation frequencies substantially in several experiments and is a necessary step to ensure accurate results (**Fig. 1B and C**, **Supplementary Fig. 1**)

Estimating mutation efficiency starts with the alignment of the sequenced reads (**Fig. 1A**). A common strategy is to use standard genomic alignment tools. However, these tools do not align using knowledge about the known mechanisms of CRISPR-induced double stranded breaks and DNA repair. Genome editing typically results in a single deletion and/or insertion of variable length. Hence, correctly aligned reads will often have a low number of events (optimally 1 deletion and/or 1 insertion after normalization for controls) overlapping the cut site, while misaligned reads will result in a high number of events throughout the read due to discrepancies to the correct loci. Therefore an alignment strategy that penalizes multiple indel events (see Methods) is more consistent with DNA repair mechanisms and the CRISPR mode of action. ampliCan uses the Needleman-Wunsch algorithm with tuned parameters to ensure optimal alignments of the reads to their loci and models the number of indel and mismatch events to ensure that the reads originated from that loci (see Methods and **Supplementary Note 2**). In contrast, non-optimized aligners can create fragmented alignments resulting in misleading mutation profiles and possible distortion of downstream analyses and frameshift estimation (**Supplementary Fig. 2**). In assessments, ampliCan outperforms the tools CrispRVariants, CRISPResso and AmpliconDivider on the synthetic benchmarking previously used to assess these tools (Lindsay et al. 2016), where experiments were contaminated with simulated off-target reads that resemble the real on-target reads, but have a mismatch rate of 30% per bp (**Supplementary Fig. 4**). A cause for concern is that the mapping strategy used in the pipelines of several tools (**Supplementary Table 1**) are not robust to small perturbations of this mismatch rate and when we simulated contaminant off-target data with varying degrees of mismatches to the on-target loci (see **Supplementary Note 4**) it lead to a significant reduction in performance (**Fig. 2**, left). In contrast, ampliCan’s strategy of modelling editing events to ascertain whether a read originated from the on-target or the off-target loci resulted in consistently high performance across a broad range of mismatch rates (**Fig. 2**, left and **Supplementary Figs. 4**, **5**).

**Fig 2.**
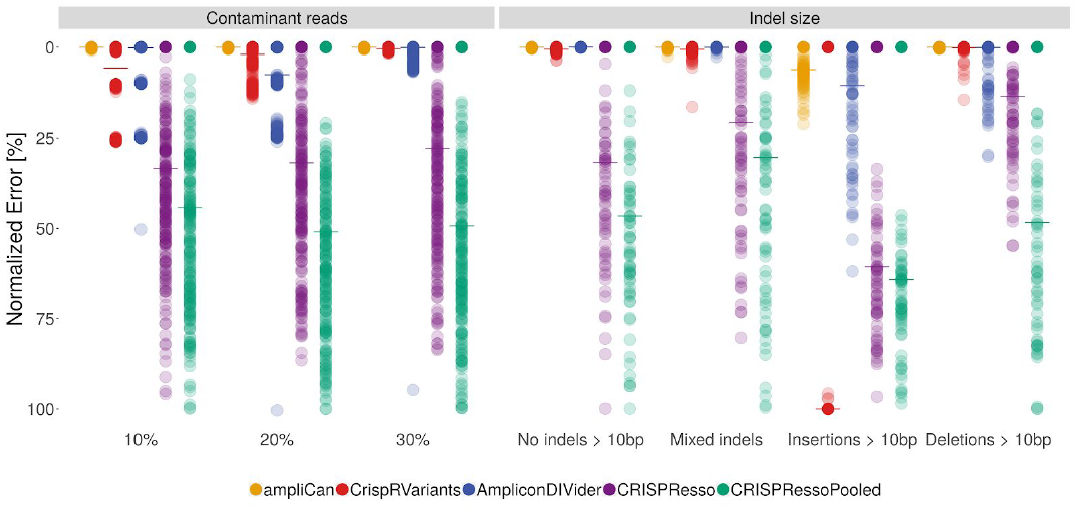
Benchmark of leading tools when estimating mutation efficiency under different dataset conditions. Each dot shows the error of the estimate to the correct value for a single experiment normalized to 0-100 scale. The median performance (Mixed indels) is indicated by the horizontal line. The left panel shows comparison of tools when datasets contain contaminant reads (see text and methods). The x-axis denotes how dissimilar the contaminant reads are to the correct reads. In cases where the contaminants are from homologous regions this may be low (10%), for other contaminants this is likely to be higher (30%). The right panel shows performance of tools as a function of the length of indel events. The sets in the left column contain no indels > 10bp, the right column contain only insertions > 10bp, the middle column (Mixed Indels) is a mix of shorter and longer events.

### ampliCan can detect long indels and estimate HDR efficiency

Targeted insertion of shorter fragments through co-opting of the homology directed repair (HDR) pathway is becoming increasingly popular (Kuscu et al. 2017; Lackner et al. 2015). This, together with long indels occurring in regular CRISPR experiments, presents a challenge for most CRISPR analysis tools. To assess the ability of the leading tools in recognizing long indels we simulated data using the strategy from Lindsay et al. 2016, but restricting to indels of 10bp or longer. This revealed an inability of current pipelines to process these longer events (**Fig. 2**, right) typically stemming from alignment strategies that are unable to assign reads with long indels to the correct loci. In previous assessments, simulated data has often been restricted to short indels where this weakness would not be apparent (**Supplementary Note 5**). Using a localized alignment strategy, based on primer matching (see Methods), ampliCan knows *a priori* which loci the reads are supposed to originate from. This alignment strategy therefore outperforms all other tools and robustly handles these longer indels (>10bp) when they occur unintentionally (**Fig. 2**, right and **Supplementary Fig. 6**).

Intentional introduction of specific edits using a donor templates is supported in ampliCan through an HDR mode where it first aligns the donor template to the reference in order to identify editing events that are expected to take place in a successful integration. The presence of these success-events are then quantified in the edited samples obtaining the frequency of integration. To assess this strategy we simulated experiments with different levels of donor integration (a result of HDR) in the presence of different levels of cut loci but with donor introduction (a result of NHEJ). This revealed that only ampliCan can consistently recover both the true HDR and NHEJ efficiency (**Supplementary Note 6** and **Supplementary Fig. 7**). An identical strategy also makes it possible to quantify the efficiency of base editors (Gaudelli et al. 2017; Komor et al. 2016) by supplying ampliCan with templates where the target bases have been altered.

### ampliCan summarizes and aggregates results over thousands of experiments

To aid analysis of heterogeneous outcomes, ampliCan quantifies the heterogeneity of reads (**Supplementary Fig. 9**), the complete mutation efficiency for an experiment and the proportion of mutations resulting in a frameshift (**Fig. 1C**, top right). It also aggregates and quantifies mutation events of a specific type if a particular outcome is desired (**Supplementary Fig. 8**). In addition, ampliCan provides overviews of the impact of all filtering steps (**Supplementary Fig. 10**). Reports can be generated in several formats (**Supplementary Tables 2 and 3**) and aggregated at multiple levels such as sequencing barcodes, gRNA, gene, loci or any user-specified grouping (**Supplementary Note 7**). This enables exploration of questions beyond mutation efficiency such as the rules of gRNA design, whether a particular researcher is better at designing gRNAs than others (**Supplementary Fig. 11**), whether a given barcode is not working or determining the stochasticity in the mutation outcome from a given gRNA (**Supplementary Fig. 22**).

ampliCan offers a complete pipeline controlling for biases at every step of evaluation. It can be integrated with the CHOPCHOP tool for gRNA design to incorporate all computational steps necessary for a CRISPR experiment. It scales from a single gRNA to genome-wide screens and can be run with a single command. For more advanced users, it provides a complete and adaptable framework, implemented in R and bioconductor, enabling further exploration of the data. Collectively, these advances will minimize misinterpretation of genome editing experiments and allow effective analysis of the outcome in an automated fashion.

## Methods

### ampliCan pipeline

ampliCan is completely automated and accepts a configuration file describing the experiment(s) and FASTQ files of sequenced reads as input. The configuration file contains information about barcodes, gRNAs, forward and reverse primers, amplicons and paths to corresponding FASTQ files (**Supplementary Table 4**). From here, ampliCan generates reports summarizing the key features of the experiments.

In the first step, ampliCan filters low quality reads which either have ambiguous nucleotides, an average quality or individual base quality under a default or user-specified threshold (**Supplementary Note 8**). After quality filtering, ampliCan assigns reads to the particular experiment by searching for matching primers (default up to two mismatches, but ampliCan supports different stringency, **Supplementary Note 8**). Unassigned reads are summarized and reported separately for troubleshooting. After read assignment ampliCan uses the Biostrings (Pages et al.) implementation of the Needleman-Wunsch algorithm with optimized parameters (gap opening = -25, gap extension = 0, match = 5, mismatch = -4, no end gap penalty) to align all assigned reads to the loci/amplicon sequence. Subsequently, primer dimer reads are removed by detecting deletions larger than the size of the amplicon subtracting the length of the two primers and a short buffer. Additionally, sequences that contain a high number of indels or mismatch events compared to the remainder of the reads are filtered as these are potential sequencing artifacts or originate from off-target amplification (**Supplementary Note 8** and **Supplementary Fig. 13**). Mutation frequencies are calculated from the remaining reads using the frequency of indels that (**Supplementary Fig. 8**) overlap a region (+/-5bp) around the expected cut site. If paired-end sequencing is used, ampliCan follows consensus rules for the paired forward and reverse read generally picking the read with the best alignment in case of disagreement (described in **Supplementary Fig. 14**). The alternative strategy of merging the paired reads is supported by ampliCan, but has been demonstrated to be detrimental to performance (Lindsay et al. 2016). The expected cut site can be specified as a larger region for nickase or TALEN experiments where the exact site is not known. Any indel or mismatch also observed above a 1% threshold in the control are removed. Frameshifts are identified by summing the impact of deletions and insertions on the amplicon.

A series of automated reports is prepared in form of “.Rmd” files which can be converted to multiple formats, but also immediately transformed into html reports with knitr (Xie 2013) for convenience. There are six different default reports prepared by ampliCan with statistics grouped at the corresponding level: id, barcode, gRNA, amplicon, summary and group (user specified, typically person conducting experiments, treatment or other grouping of interest). In addition to alignments of top reads (**Fig. 1C, Supplementary Fig. 1**) reports contain plots summarized over all deletions, insertions and variants (**Supplementary Fig. 8**). In addition a number of plots showing the general state of the experiments are shown including the heterogeneity of reads to investigate mosaicism or sequencing issues (**Supplementary Figs. 9**, **15**, **16**) and overviews of how many reads were filtered/assigned at each step (**Supplementary Fig. 17**). In addition to the default plots ampliCan produces R objects that contain all alignments and read information, these can be manipulated, extended and visualized through the R statistical package.

ampliCan provides a versatile tool that can be used out-of-the-box or as a highly flexible framework that can be extended to more complex analysis. The default pipeline consistst of a single convenient wrapper, amplicanPipeline, which generates all default reports. More advanced users can gain complete control over all processing steps (**Supplementary Fig. 12**) and produce novel plots for more specialized use cases. Compatibility with the most popular plotting packages ggplot2 (Wickham and Wickham 2007) and ggbio (Yin et al. 2012) as well as the most popular data processing packages dplyr (Wickham and Francois 2015) and data.table provides a full fledged and elastic framework. Output files are encoded as GenomicRanges (Lawrence et al. 2013) tables of aligned read events for easy parsing (**Supplementary Table 3**) and human readable alignment results (**Supplementary Table 2**) and fasta. We would like to encourage users to communicate their needs and give us feedback, for future development.

### Running parameters

All the tools were used with their default options, specific versions of the tools and software is specified in the https://github.com/valenlab/amplican_manuscript description file.

## Data access

### Data Availability

All real datasets come from zebrafish TLAB strain and are available online under accession numbers: PRJNA245510 (BioProject, run 1 and run 5) and E-MTAB-6310, E-MTAB-6355, E-MTAB-6356, E-MTAB-6357, E-MTAB-6358, (run 6-10). Descriptions, treatments and other details of those datasets are described in the (Gagnon et al. 2014). Real datasets were used to analyze influence of usage of control on editing efficiency estimations and prove that the tools estimations of efficiency estimates differ (**Supplementary Fig. 3**). Synthetic datasets can be reconstructed with the use of code from https://github.com/valenlab/amplican_manuscript. Synthetic datasets were created in a similar fashion to the sets in (Lindsay et al. 2016) using 20 different loci edited at variable efficiency (0, 33.3, 66.7 and 90%) and with the possibility of adding HDR. Further details can be found in the **Supplementary Data Overview**.

### Code availability

ampliCan is developed as an R package under GNU General Public License version 3 and available through Bioconductor under http://bioconductor.org/packages/amplican or https://github.com/valenlab/amplican. Automatic HDR estimation is available since version 1.1.4, improved HDR version is available from version 1.3.3.

## Acknowledgements

We would like to thank Jason Rihel, Tessa Montague and Alex Schier for support and many useful comments, and members of the Schier lab for their contributions.

The project was supported by the Bergen Research Foundation and the Norwegian Research Council (FRIMEDBIO #250049) (E.V.), University of Bergen core funding (K.L.) and the American Cancer Society and University of Utah startup funding (J.A.G.).

## Author Contributions

E.V. conceived and supervised the project. J.A.G. performed wet-lab experiments and prepared datasets. X.G., G.C. and A.C. assisted in data interpretation and writing the manuscript. K.L. developed the R package and performed all computational work.

## References

Cong L, Ran FA, Cox D, Lin S, Barretto R, Habib N, Hsu PD, Wu X, Jiang W, Marraffini LA, et al. 2013. Multiplex genome engineering using CRISPR/Cas systems. Science 339: 819–823.

Courtney DG, Moore JE, Atkinson SD, Maurizi E, Allen EHA, Pedrioli DML, McLean WHI, Nesbit MA, Moore CBT. 2016. CRISPR/Cas9 DNA cleavage at SNP-derived PAM enables both in vitro and in vivo KRT12 mutation-specific targeting. Gene Ther 23: 108–112.

Gagnon JA, Valen E, Thyme SB, Huang P, Akhmetova L, Ahkmetova L, Pauli A, Montague TG, Zimmerman S, Richter C, et al. 2014. Efficient mutagenesis by Cas9 protein-mediated oligonucleotide insertion and large-scale assessment of single-guide RNAs. PLoS One 9: e98186.

Gaudelli NM, Komor AC, Rees HA, Packer MS, Badran AH, Bryson DI, Liu DR. 2017. Programmable base editing of A•T to G•C in genomic DNA without DNA cleavage. Nature 551: 464.

Jinek M, Chylinski K, Fonfara I, Hauer M, Doudna JA, Charpentier E. 2012. A programmable dual-RNA-guided DNA endonuclease in adaptive bacterial immunity. Science 337: 816–821.

Komor AC, Kim YB, Packer MS, Zuris JA, Liu DR. 2016. Programmable editing of a target base in genomic DNA without double-stranded DNA cleavage. Nature 533: 420–424.

Kuscu C, Parlak M, Tufan T, Yang J, Szlachta K, Wei X, Mammadov R, Adli M. 2017. CRISPR-STOP: gene silencing through base-editing-induced nonsense mutations. Nat Methods 14: 710–712.

Labun K, Montague TG, Gagnon JA, Thyme SB, Valen E. 2016. CHOPCHOP v2: a web tool for the next generation of CRISPR genome engineering. Nucleic Acids Res 44: W272–6.

Lackner DH, Carré A, Guzzardo PM, Banning C, Mangena R, Henley T, Oberndorfer S, Gapp BV, Nijman SMB, Brummelkamp TR, et al. 2015. A generic strategy for CRISPR-Cas9-mediated gene tagging. Nat Commun 6: 10237.

Lawrence M, Huber W, Pagès H, Aboyoun P, Carlson M, Gentleman R, Morgan MT, Carey VJ. 2013. Software for computing and annotating genomic ranges. PLoS Comput Biol 9: e1003118.

Lindsay H, Burger A, Biyong B, Felker A, Hess C, Zaugg J, Chiavacci E, Anders C, Jinek M, Mosimann C, et al. 2016. CrispRVariants charts the mutation spectrum of genome engineering experiments. Nat Biotechnol 34: 701–702.

Montague TG, Cruz JM, Gagnon JA, Church GM, Valen E. 2014. CHOPCHOP: a CRISPR/Cas9 and TALEN web tool for genome editing. Nucleic Acids Res 42: W401–7.

Pages H, Gentleman R, Aboyoun P, DebRoy S. Biostrings: String objects representing biological sequences, and matching algorithms, 2008. R package version 2: 160.

Wickham H, Francois R. 2015. dplyr: A grammar of data manipulation. R package version 0 4 1: 20.

Wickham H, Wickham MH. 2007. The ggplot package. http://ftp.uni-bayreuth.de/math/statlib/R/CRAN/doc/packages/ggplot.pdf.

Xie Y. 2013. knitr: A general-purpose package for dynamic report generation in R. R package version 1: 1.

Yin T, Cook D, Lawrence M. 2012. ggbio: an R package for extending the grammar of graphics for genomic data. Genome Biol 13: R77.

